# Inhibition of *Pseudomonas aeruginosa* quorum sensing by chemical induction of the MexEF-oprN efflux pump

**DOI:** 10.1101/2023.10.27.564458

**Authors:** Rasmus Kristensen, Jens Bo Andersen, Morten Rybtke, Charlotte Uldahl Jansen, Blaine Gabriel Fritz, Rikke Overgaard Kiilerich, Jesper Uhd, Thomas Bjarnsholt, Katrine Qvortrup, Tim Tolker-Nielsen, Michael Givskov, Tim Holm Jakobsen

**Author notes:** Corresponding author: Tim Holm Jakobsen, Costerton Biofilm Center, Department of Immunology and Microbiology, Faculty of Health and Medical Sciences, University of Copenhagen. Blegdamsvej 3B, DK-2200 Copenhagen N, Denmark.

## Abstract

The cell-to-cell communication system quorum sensing (QS), used by various pathogenic bacteria to synchronize gene expression and increase host invasion potentials, is studied as a potential target for persistent infection control. To search for novel molecules targeting the QS system in the Gram-negative opportunistic pathogen *Pseudomonas aeruginosa* a chemical library consisting of 3280 small compounds from LifeArc was screened. A series of ten conjugated phenones that have not previously been reported to target bacteria were identified as inhibitors of QS in *P. aeruginosa*. Two lead compounds (ethylthio enynone and propylthio enynone) were resynthesized for verification of activity and further elucidation of the mode of action. The isomeric pure Z-ethylthio enynone was used for RNA sequencing revealing a strong inhibitor of QS-regulated genes and the QS-regulated virulence factors rhamnolipid and pyocyanin were significantly decreased by treatment with the compounds. A transposon mutagenesis screen performed in a newly constructed *lasB*-*gfp* monitor strain identified the target of Z-ethylthio enynone in *P. aeruginosa* to be the MexEF-OprN efflux pump, which was further established using defined *mex* knockout mutants. Our data indicate that the QS inhibitory capabilities of Z-ethylthio enynone were caused by the drainage of intracellular signal molecules as a response to chemical-induced stimulation of the MexEF-oprN efflux pump thereby inhibiting the auto-generated positive feedback and its enhanced signal-molecule synthesis.

## INTRODUCTION

*Pseudomonas aeruginosa* has been the subject of extensive research and development efforts aimed at combating its persistent infections, which have been recognized as a high priority by the World Health Organization (1). The large genome of *P. aeruginosa* contains a plethora of metabolic and virulence factor-related genes, making this bacterium highly adaptive and versatile when it comes to living in harsh environments (2, 3). While *P. aeruginosa* is typically isolated from soil, it can utilize its versatility to act as an opportunistic pathogen by colonizing patients, most notoriously in the lower respiratory tract of cystic fibrosis patients (4), as well as chronic wounds (5) and at various body sites during nosocomial infections (6).

Host colonization relies on a well-coordinated synthesis of virulence factors on a community level to ensure the host invasion is successful. *P. aeruginosa* achieves this by using cell-to-cell communication known as quorum sensing (QS), which utilizes chemical signaling to ensure regulation of the pathogenicity in a cell-density-dependent manner. Discovered in the bacterium *Vibrio fischeri,* the biosynthesis of the signal molecule *N*-acyl homoserine lactone (AHL) and its interaction with the *luxI-luxR* system gave insight into how bacteria can “communicate” by simple diffusion (7). The signal molecule concentration produced by the individual cell is insufficient to self-activate the system, but as the community size increases, likewise, the concentration of the surrounding signal molecules until the QS activation threshold has been reached, leading to upregulation of QS-related genes. *P. aeruginosa* possesses two *luxI-luxR* homologous systems working in a hierarchical order, firstly *lasI-lasR* and secondly *rhlI-rhlR*, each consisting of a signal synthetase lasI/rhlI and receptor proteins lasR/rhlR for initiating transcription (8). The main differences between these two systems are the chemical structure of their signal molecules, with the Las system synthesizing and reacting to n-3-oxo-dodecanoyl-l-homoserine lactone (3-oxo-C12-HSL) (9) and the Rhl system to N-butyryl-l-homoserine lactone (C4-HSL) (10), thereby circumventing cross system interference. With the binding of 3-oxo-C12-HSL to LasR, a positive feedback loop is created, resulting in increased synthesis of 3-oxo-C12-HSL as well as initiation of the production of C4-HSL by *rhlI* (11). Additional regulation of the QS systems occurs through the pseudomonas quinolone signal (*PQS*), in which conversion from a precursor molecule to 2-heptyl-3-hydroxy-4-quinolone (PQS) is dependent on the expression of *lasR* (12). While knockout mutants lacking parts of the *PQS* gene cluster have greatly reduced synthesis of virulence factors associated with the Rhl system, the exact mechanism, which connects PQS to the Rhl system has not been fully outlined (13). Despite this, it is widely accepted that PQS is a primary mediator between the Las and Rhl system. It is estimated that up to 6% of the *P. aeruginosa* genome consists of QS and QS-regulated virulence genes (14). Therefore, inhibition of the QS system and pathogenicity is a potential target for the development of anti-virulence compounds.

Among the secreted virulence factors of *P. aeruginosa* are the Rhl system-regulated rhamnolipid and pyocyanin, which are two important factors in host colonization. Rhamnolipids are surfactants that facilitate bacterial motility and have cytotoxic activity towards polymorphonuclear leukocytes and macrophages leading to necrosis (15). Pyocyanin belongs to a group of secondary metabolites called phenazines known for their characteristic vibrant green/blue color found in *P. aeruginosa* infections (16). By crossing the host cell membrane and functioning as a redox-active compound, pyocyanin can generate intracellular oxidative stress using reactive oxygen species affecting the mitochondria (17, 18). This leads to fatal cell damage and elevated inflammation by recruitment of additional immune cells (19).

*P. aeruginosa* possesses intrinsic resistance mechanisms towards multiple classes of antibiotics, including the ability to export toxic molecules out of the cell by the use of multiple efflux pumps. These pumps belong to the resistance-nodulation-cell division (RND) family of which twelve have been identified in *P. aeruginosa* (20). The RND efflux pumps consist of three proteins that form a channel in the cell envelope and facilitate efflux from the cytoplasm to the surrounding environment under tight regulation at a local and global level (21). While elevated activity of the efflux pumps correlates with appearance of antibiotic resistant phenotypes, it is broadly recognized that the primary functions of these pumps are elements of an adaptive stress response towards harmful products, either metabolic byproducts or foreign elements (22). The efflux pump MexEF-OprN is unique among the RND systems found in *P. aeruginosa*, by being regulated by the activator MexT. The MexT protein is a LysR-type transcription factor regulated by toxic secondary metabolites acting as affinity-creating co-inducers that causes the transcription factor to bind to DNA and initiate gene expression (23). The concentration of the co-inducers is believed to be reduced by the quinone oxidoreductase MexS that has the ability to detoxify the MexT-activating molecules (24). Mutations in the *mexS* gene results in greater resistance toward chloramphenicol and quinolones (25). The elevated antibiotic resistance originates from the buildup of toxic stress which can only be relieved by continuous efflux created by upregulated *mexT* and increased MexEF-OprN activity. The PQS precursor HHQ has been shown to be exported out of the cell by the pump (26) which means that *mexS* mutants are less virulent as the intracellular concentrations of the QS signal molecules fail to reach its activation threshold (27, 28).

In the present study, a compound library provided by LifeArc (formerly MRC Technology, London, UK) was screened for QS inhibitory activity. Ten compounds were selected for further analysis based on inhibitory activity, and two compounds (ethylthio enynone and propylthio enynone) were re-synthesized for hit validation. The isomeric pure Z-ethylthio enynone was employed in target identification studies. Utilizing mariner transposon mutagenesis assay, it was determined that the QS inhibitory activity was caused by the activation of the MexEF-OprN efflux pump.

## RESULTS

### Library screening for compounds capable of decreasing expression of QS-regulated genes

A library consisting of 3280 compounds was screened for inhibitory activity of the expression of the *P. aeruginosa* QS-regulated gene, *rhlA.* For the entire screening and the subsequent identification of hit compounds, cell-based systems employing fluorescence as an indicator of gene expression of selected QS-regulated genes were used. The systems are based on an unstable version of green fluorescent protein (GFP(ASV)) (29) fused to the promoter of a QS-regulated gene (*lasB* or *rhlA*) in *P. aeruginosa* PAO1 (**Table S1**). The library provided by LifeArc consisted of primarily natural product compounds divided into two series with 1200 and 2080 compounds. In the initial screen, the potency of the compounds at a concentration of 10 µM was assessed by measuring fluorescence at a single time point after 12 hours of growth. Growth was measured simultaneously, and fluorescence measurements were normalized if any changes in growth were observed. From the initial screen, 89 compounds were found to decrease the expression of *rhlA* by more than 20% when compared to an untreated control (**Fig. 1**). From a secondary test of the 89 compounds in serial dilutions with both the *rhlA-gfp* (30) and the *lasB-gfp* (31) monitor strains 10 compounds were selected, and new batches were tested in 10 times higher concentrations in order to validate and calculate meaningful IC_50_ values. The molecular structures of the compounds were provided by LifeArc giving us the opportunity to consider the novelty value and relevance to further investigate the selected compounds. To our knowledge, none of the selected compounds have been investigated previously for targeting *P. aeruginosa*. The molecular structures consist of a phenone moiety conjugated to an alkyne followed by either an alkene (946, 950, 952, 953, 955, 963 and 964, **Fig. 1**) or an alkyne (944 and 959, **Fig. 1**). For the majority of the compounds the alkene is in conjugation with an alkylthiol. The chemical structures of the identified compounds are different from previously investigated compounds targeting the QS signaling system. Therefore, our hypothesis was that these compounds may possess a novel target, thus far unexplored in studies investigating small compounds that can attenuate the virulence of *P. aeruginosa*.

**FIG 1.**
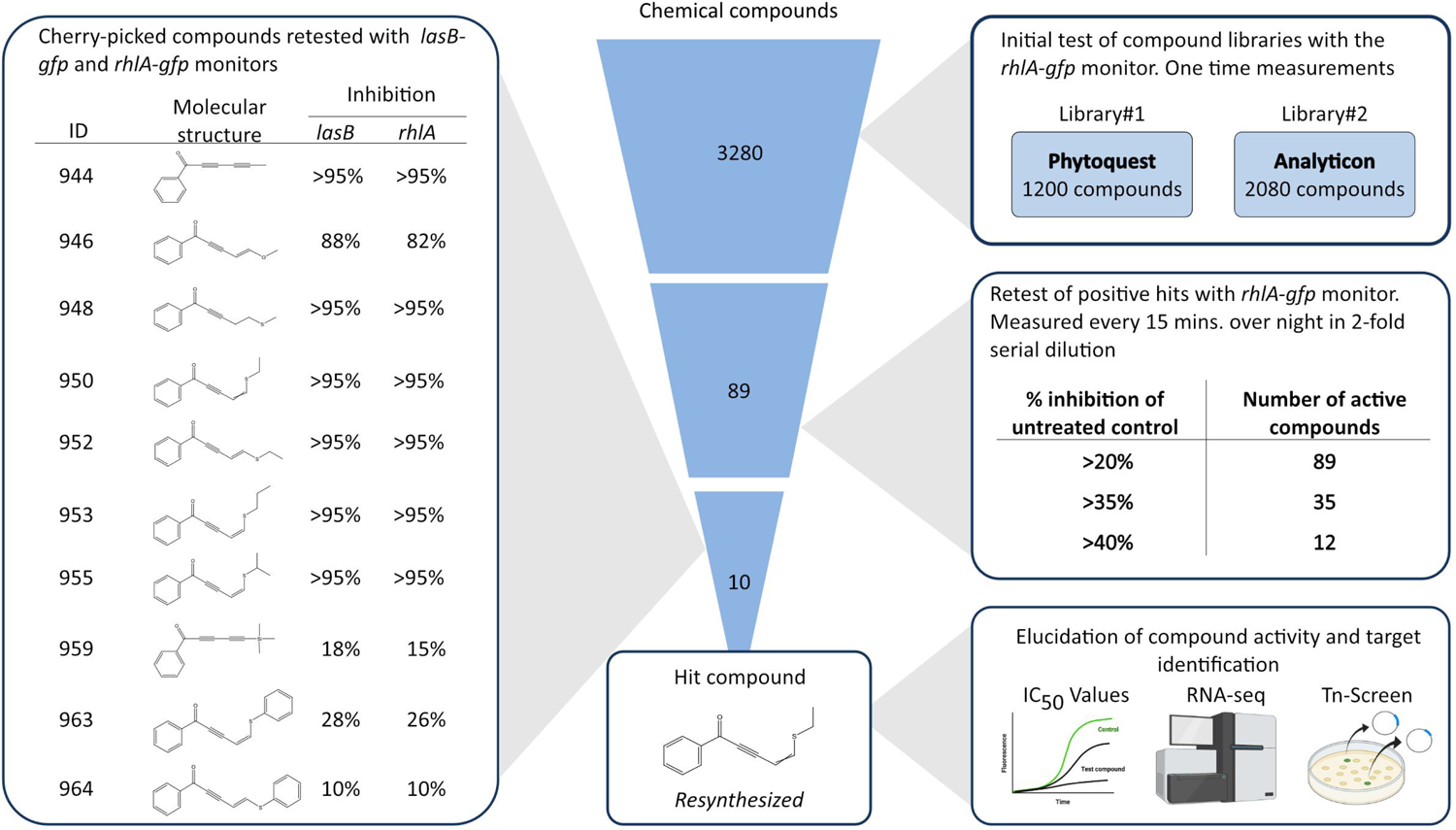
Compound library screen. From the 3280 compounds tested for quorum sensing inhibition with the *rhlA-gfp* monitor 89 compounds were found to decrease the fluorescence output more than 20%. The 89 compounds were retested and divided according to the percentage inhibition of fluorescence in a 10 μM concentration. From the retest, 10 compounds were selected, and new batches were tested in 2-fold serial dilutions from 100 μM concentrations with the *rhlA-gfp* and *lasB-gfp* monitor strains. Two hit compounds were resynthesized and selected for further analysis and target identification.

### Synthesis of hit compounds

The selected compounds were synthesized by addition of a mono-TMS-protected diyne to benzaldehyde, followed by an oxidation with manganese dioxide to form the ketone. Lastly, selective nucleophilic addition of the thiol to the terminal triple bond formed the desired alkylthio enynone. The synthetic route can be seen in **Fig. 2**.

**FIG 2.**
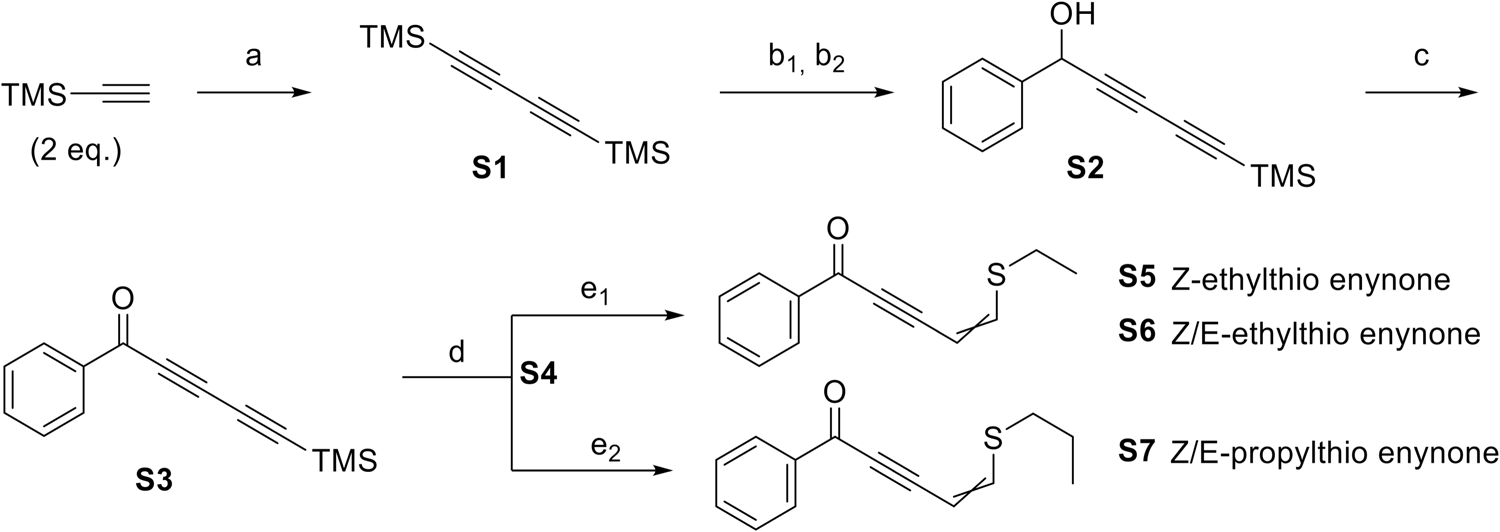
Synthetic route: (a) CuI (0.09 eq.), TMEDA (0.21 eq.), DIPEA (3 eq.), THF (3.9 mL/mmol alkyne), 60°C overnight. (b_1_) MeLi·LiBr (1.5M, 1 eq.), THF (3.85 mL/mmol **S1**), 0°C→rt. 4 hours. (b_2_) benzaldehyde (1.2 eq.), 0°C 1 hour. (c) MnO_2_ (20 eq.), DCM (5.02 mL/mmol **S2**), reflux 2.5 hours. (d) Na_2_CO_3_:SiO_2_ (1:1), EtOAc (24 mL/mmol **S3**), rt. 11 hours. (e_1_) EtSH (1.1 eq.), (n-Bu)_3_P (0.1 eq.), DCM (7 mL/mmol **S4**), rt. 1 minuet. (e_2_) PrSH (1.1 eq.), (n-Bu)_3_P (0.1 eq.), DCM (7 mL/mmol **S4**), rt. 1 minuet.

The dual TMS protected diyne (**S1**) was synthesized by a Hay homo-coupling of TMS-acetylene, followed by selective mono-deprotection using one equivalent of the MeLi·LiBr complex at low temperature. Addition of benzaldehyde to the reaction gave the secondary alcohol (**S2**) (32), which was afterwards oxidized to give the ketone (**S3**) (32). Removal of the TMS group NaCO_3_:SiO_2_, allowed addition of the appropriate alkylthiol to the alkyne (**S4**) using (n-Bu)_3_P as base to afford the alkylthio enynone products (**S6** or **S7**). The synthesized compounds were purified by preparative HPLC, allowing isolation of the pure Z-isomer of ethylthio enynone (**S5**) as well as an isomeric mixture of the Z/E-ethylthio enynone isomeric mixture (**S6**). For the propylthio enynone (**S7**) it was not possible to separate the two isomers. Synthetic procedures can be found in Supplemental Material.

### Expression of selected QS-regulated genes lowered by alkylthio enynone compounds

The isomeric pure Z-ethylthio enynone (**S5**), as well as the ethylthio enynone isomeric mixture (**S6)**, and the propylthio enynone isomeric mixture (**S7**), (**Fig. 3A**) were all used for additional testing.

**FIG 3.**
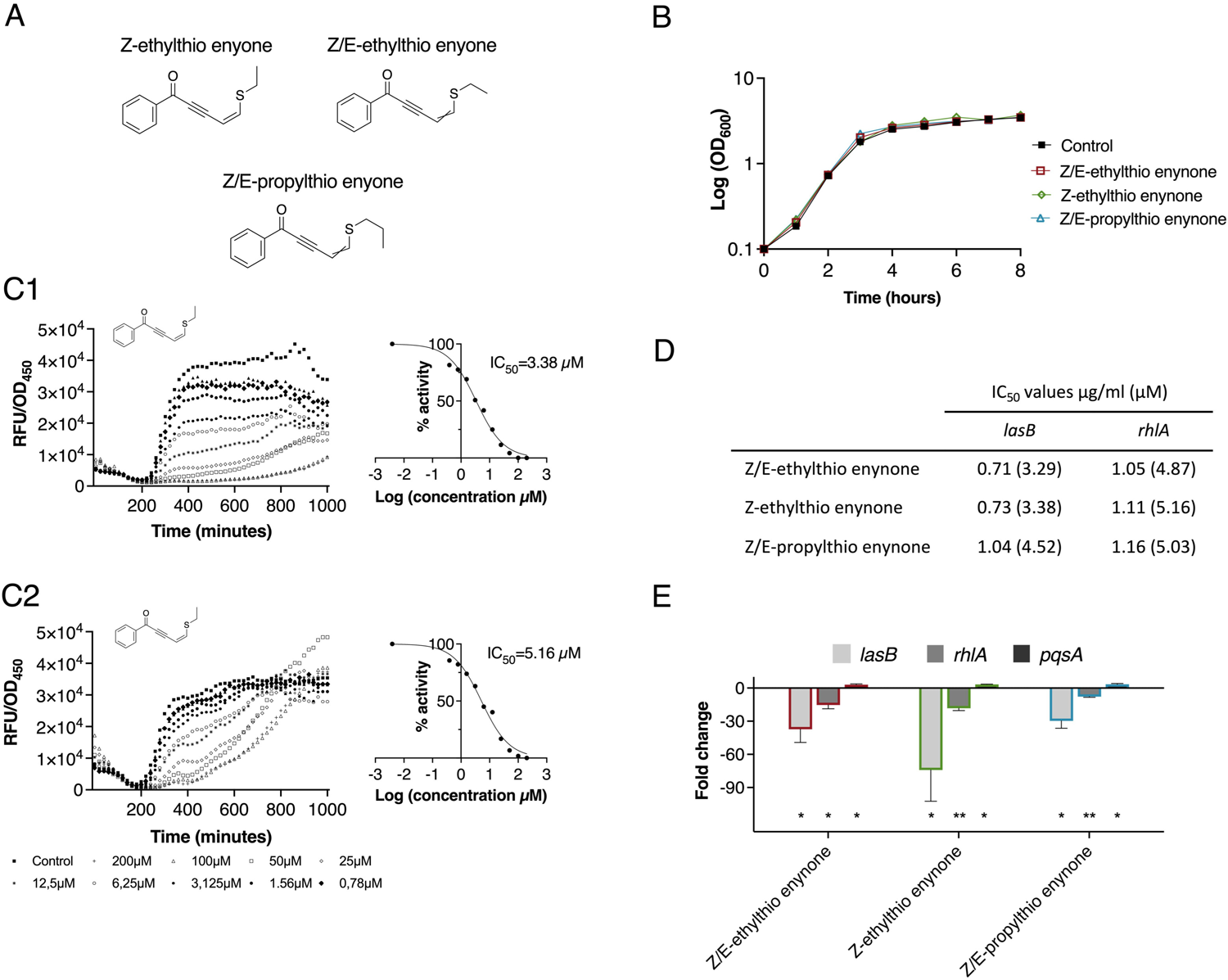
Inhibitory potency of resynthesized alkylthio enynone compounds. A) Chemical structures of synthesized compounds investigated, Z-ethylthio enynone, Z/E-ethylthio enynone and Z/E-propylthio enynone. B) Growth measured as cell density (OD_600_) of untreated and treated WT PAO1 with 100 μM of the investigated compounds. C) Dose-response curves (specific fluorescence) of Z-ethylthio enynone incubated with the monitor *lasB-gfp* (C1) and *rhlA-gfp* (C2) with corresponding curves for calculation of IC_50_ values. D) IC_50_s of investigated compounds calculated using the *lasB-gfp* and *rhlA-gfp* monitor strains. E) Fold change in gene expression of *lasB*, *rhlA* and *pqsA* in WT PAO1 grown with 100 μM of Z/E-ethylthio enynone, Z-ethylthio enynone and Z/E-propylthio enynone measured by quantitative real-time PCR. The results are based on three independent experiments. Error bars represent means ± standard deviations (SD). Significance levels from zero are based on one sample t-test analysis (* p ≤ 0.05 and ** p ≤ 0.01).

None of the compounds showed growth inhibitory activity in 100 μM concentrations (**Fig. 3B**). The Z-isomer and E/Z-isomer of ethylthio enynone lowered the expression of *lasB* and *rhlA* in a very similar fashion with fifty percent inhibitory concentrations (IC_50_s) values of approximately 3.3 μM (0.7 μg/ml) and 5 μM (1 μg/ml), respectively **(Fig. 3D)**, suggesting high similarity in the activity of the Z-isomer and E-isomer. The Z/E-propylthio enynone showed a slightly higher IC_50_ value when tested with the *lasB-gfp* monitor 4.52 μM (1.04 μg/ml) and very similar IC_50_ value for inhibition of the *rhlA* (5 μM (1.16 μg/ml)) as the Z-isomer and the Z/E-isomer of ethylthio enynone. The IC_50_ values were calculated from the slope which represents the synthesis rate of the dose-response curves expressing the specific fluorescence (GFP expression/cell density) shown in **Fig. 3C1+2** and **Fig. S1+S2**. To further investigate the activity of the alkylthio enynone compounds on expression of the QS-regulated *lasB, rhlA* and *pqsA* genes quantitative real-time PCR (qRT-PCR) analysis was performed. Exponentially growing *P. aeruginosa* cultures were treated with each of the compounds in concentrations of 100 μM. RNA was extracted from samples retrieved at an OD of 2.0 (stationary growth phase). Previous studies have reported that at this point in the growth phase, the majority of the QS-regulated genes are expressed at the highest level (33). The qRT-PCR data showed that the three test compounds caused a significant decrease in transcription of *lasB* and *rhlA*, whereas there was a 3-fold increase in *pqsA* transcription. The Z/E-ethylthio enynone and the Z-ethylthio enynone showed the highest reduction in *lasB* expression (37-fold and 74-fold, respectively) compared to *rhlA* (15-fold and 18-fold, respectively), while the Z/E-propylthio enynone showed the lowest inhibitory effect on *lasB* (30-fold) and on *rhlA* (8-fold) (**Fig. 3E**). To test whether the three compounds affect the expression of the QS genes through the Gac/Rsm cascade we used the two monitor strains *rsmY-gfp* and *rsmZ-gfp* (34). None of the compounds changed the fluorescence signal of these monitor strains, which indicates that the compounds do not target the Gac/Rsm cascade (**Fig. S4**). The results of the experiments with the five different monitor strains and the qRT-PCR analysis indicate that the compound specifically targets the Las and Rhl part of the QS system, whereas, it has no or a very modest positive effect on the PQS part of the QS system.

### Multiple QS-regulated virulence mechanisms are targeted by Z-ethylthio enynone treatment in *P. aeruginos*a

To further investigate the effect of alkylthio enynone on the *P. aeruginosa* transcriptome we employed RNA sequencing. By gaining insight into target gene specificity, we anticipated that indications of a potential target could be revealed for further investigations. Although all three hit compounds demonstrated similar QSI activity, Z-ethylthio enyone was chosen as the leading compound for downstream experiments focusing on identifying the mechanism of action, as the synthesis yield of this particular compound was the highest among the three tested compounds included in this study. Exponentially growing *P. aeruginosa* batch cultures were treated with 100 μM Z-ethylthio enynone and samples were retrieved at OD_600_ of 2.0. The purified RNA was used for the qRT-PCR analysis presented in **Fig. 3** and the RNA-seq analysis. In total 244 genes were significantly (*P*<0.05) downregulated more than 5-fold by the Z-ethylthio enynone compound, and of those 91 were QS-regulated genes according to the QS regulon defined by Hentzer et al. (35) **(Fig. 4A).** Of the significantly upregulated genes (>5 fold) 3 were QS regulated of a total of 136 genes. A high number of QS-regulated genes were downregulated, however, the specificity is relatively low (38%) suggesting that the compound is not targeting the QS system directly. Approximately 65% of the genes that were significantly changed were downregulated. A large number of the downregulated genes belong to the classification “Secreted factors” as these genes are responsible for the synthesis of various QS-related virulence factors such as pyocyanin and rhamnolipid. “Transport of small molecules” is among the categories with the largest number of upregulated genes, however at this stage, no evidence supported that these genes were responsible for Z-ethylthio enynone effect on the QS (**Fig. 4B**).

**FIG 4.**
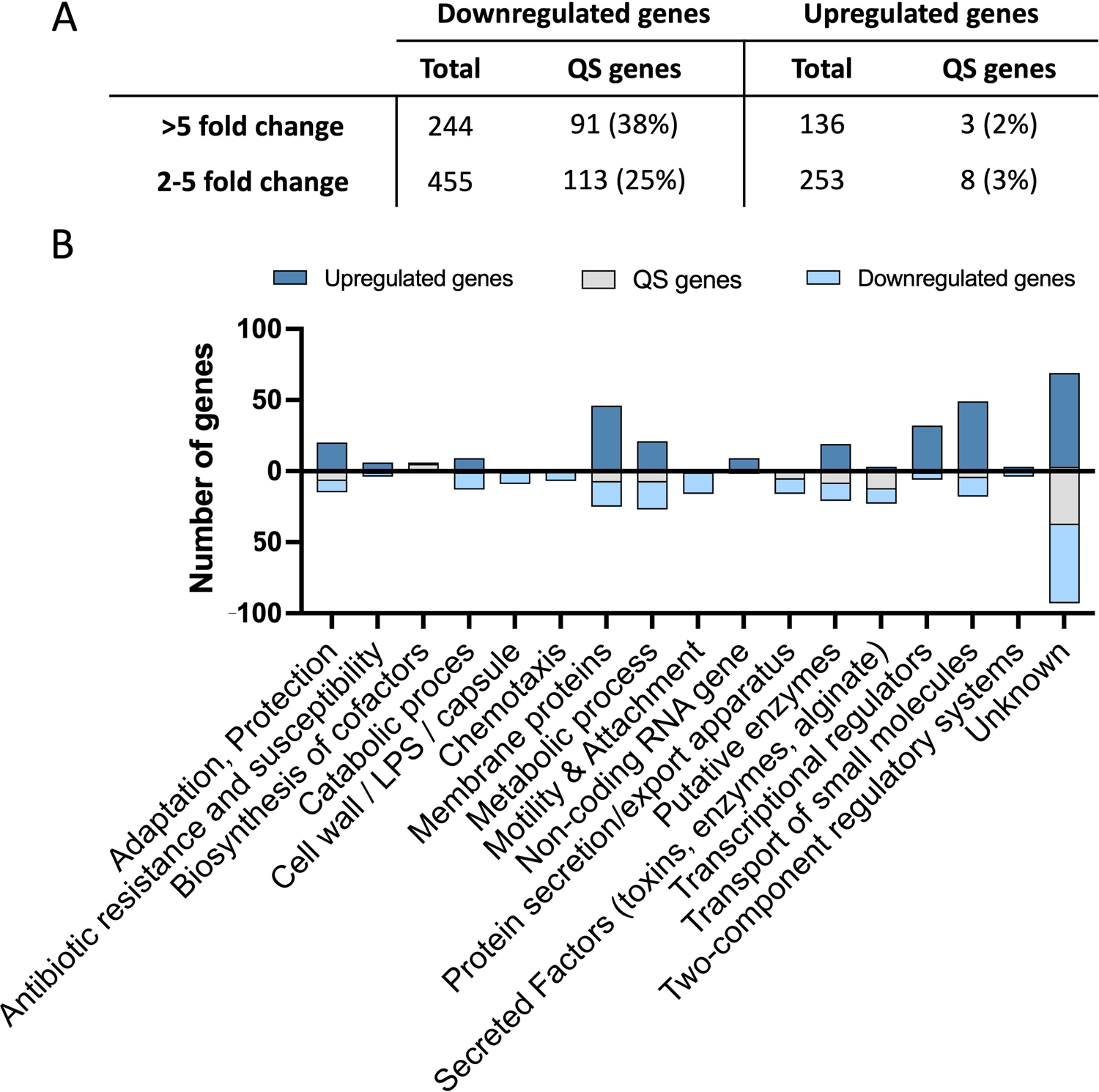
Genes significantly changed by Z-ethylthio enynone. RNA-seq analysis of PAO1 treated with 100 μM Z-ethylthio enynone. A) Total number of genes up- or down-regulated divided according to different fold changes. QS genes are genes previously identified as part of the QS regulon by (35). B) Functional classification of the top 300 up- and downregulated genes as assigned by the *P. aeruginosa* Community Annotation Project of the Pseudomonas database. Dark blue: Upregulated genes, light blue: downregulated genes, grey: genes being a part of the QS regulon according to Hentzer et al., (35).

Among the particularly interesting genes significantly affected by Z-ethylthio enynone were those encoding different virulence factors (**Table 1**), such as genes encoding rhamnolipid production, *rhlA* (PA3479) and *rhlB* (PA3478), *lasA* (1871) and *lasB* (PA3724), which are involved in the production of elastase, the two phenazine operons *phzA1-G1* (PA4210-PA4216) and *phzA2-G2* (PA1899-PA1905) encoding phenazine biosynthesis proteins, *phzM* (PA4209) encoding a putative phenazine specific methyltransferase and *phzS* (PA4217) encoding a flavin-containing monooxygenase involved in the synthesis of pyocyanin. Expression of cytogalactophillic lectin *lecA* (PA2570) and *lecB* (PA3361), which binds to oligo- and polysaccharides located on human cells (33) thereby acting as adhesion factors (31), were also affected by Z-ethylthio enynone. The precise role of LecA and LecB in *P. aeruginosa* biofilm maturation is still somewhat unclear, however, they are believed to be necessary for the early development of biofilm by forming a ligand between psl and the lectins thereby initiating biofilm formation (30). Mutations in *lecA* and *lecB* leads to more fragile and easily dispersed biofilm. Expression of the gene *cbpD* (PA0852), encoding a chitin oxidizing virulence factor that promotes survival in human blood by attenuation of the terminal complement cascade in human serum (36) and *chiC* (PA2300) encoding chitinase, were also affected by Z-ethylthio enynone. The expression of the central AHL mediated controllers of the QS system (*lasI, rhlI, lasR, rhlR*) were only slightly decreased by the treatment with Z-ethylthio enynone, whereas all genes of the *pqsABCDE* operon were upregulated by 3-fold. (The 50 most downregulated and upregulated genes by Z-ethylthio enynone are shown in **Table S3** and **Table S4)**.

**Table 1.** Alterations in expression of selected O5-regulated genes by 100 μM Z-ethylthio enynone^a^.

| Downregulated QS virulence genes |  |  |  |  |  |
| --- | --- | --- | --- | --- | --- |
| Gene nu. | Gene name | Pathogenesis factor | Function | Fold change | Adjusted P-value |
| PA0051 | *phzH* | Phenazines | oxidative stress | -5 | 9,28E-38 |
| PA0122 | *rahU* | Aegerolysins | Pore-forming toxin | -20 | 1,14E-75 |
| PA1148 | *toxA* | Toxic A | Cell death | -1 | 0,22 |
| PA0852 | *cbpD* |  |  | -33 | 3,93E-281 |
| PA1874 | - | Hypothetical protein (efflux) | Transport | -46 | 3,04E-71 |
| PA1899 | *phzA2* | Phenazines | oxidative stress | -39 | 5,15E-35 |
| PA1900 | *phzB2* | Phenazines | oxidative stress | -78 | 1,39E-139 |
| PA1901 | *phzC2* | Phenazines | oxidative stress | -46 | 3,04E-71 |
| PA1902 | *phzD2* | Phenazines | oxidative stress | -12 | 0,06 |
| PA1903 | *phzE2* | Phenazines | oxidative stress | -15 | 0,05 |
| PA1904 | *phzF2* | Phenazines | oxidative stress | -12 | 0,08 |
| PA1905 | *phzG2* | Phenazines | oxidative stress | -8 | 2,54E-12 |
| PA1871 | *lasA* | Protease | host immune evasion | -32 | 2,86E-184 |
| PA2193 | *hcnA* | Hydrogen cyanide | Poor lung function | -4 | 7,20E-10 |
| PA2194 | *hcnB* | Hydrogen cyanide | Poor lung function | -4 | 5,81E-11 |
| PA2195 | *hcnC* | Hydrogen cyanide | Poor lung function | -4 | 1,78E-21 |
| PA2570 | *lecA* | Lectin A | Adhesion, cytotoxic | -156 | 4,19E-86 |
| PA3361 | *lecB* | Lectin B | Adhesion | -22 | 4,41E-75 |
| PA3478 | *rhlB* | Rhamnolipid | Immune evasion,motility | -15 | 4,00E-77 |
| PA3479 | *rhlA* | Rhamnolipid | Immune evasion,motility | -17 | 1,34E-58 |
| PA3692 | *lptF* | Lipotoxin F | Resistance to oxidative stress and adhesion | -8 | 2,20E-21 |
| PA3724 | *lasB* | elastase | Iron acquisition | -76 | 7,97E-197 |
| PA4175 | *piv* | Endoprotease (protease IV) | Tissue damage | -39 | 5,15E-35 |
| PA4209 | *phzM* | Phenazines | oxidative stress | -16 | 8,54E-13 |
| PA4210 | *phzA1* | Phenazines | oxidative stress | -6 | 1,08E-09 |
| PA4111 | *phzB1* | Phenazines | oxidative stress | -26 | 4,70E-92 |
| PA4212 | *phzC1* | Phenazines | oxidative stress | -20 | 1,14E-75 |
| PA4213 | *phzD1* | Phenazines | oxidative stress | -4 | 0,38 |
| PA4214 | *phzE1* | Phenazines | oxidative stress | -3 | 0,31 |
| PA4215 | *phzF1* | Phenazines | oxidative stress | -7 | 0,33 |
| PA4216 | *phzG1* | Phenazines | oxidative stress | -8 | 2,20E-21 |
| PA4217 | *phzS* | Phenazines | oxidative stress | -4 | 3,00E-27 |
| - | *hsiAH* | T6SS (Phospholipase D) | Cell lysis |  | Table S6 |
| Upregulated QS virulence genes |  |  |  |  |  |
| PA0996 | *pqsA* |  | Virulence and rhl regulator | 3 | 1,15E-08 |
| PA0997 | *pqsB* |  | Virulence and rhl regulator | 3 | 5,11E-10 |
| PA0998 | *pqsC* |  | Virulence and rhl regulator | 3 | 8,37E-10 |
| PA0999 | *pqsD* |  | Virulence and rhl regulator | 3 | 7,96E-17 |
| PA1000 | *pqsE* |  | Pseudomonas type III repressor gene C | 3 | 8,75E-105 |
| Central QS regulators |  |  |  |  |  |
| PA1430 | *lasR* |  | Transcriptional Regulation | -3 | 1,53E-11 |
| PA1431 | *rsaL* |  | Transcriptional Regulation | -2 | 5,18E-04 |
| PA1432 | *lasI* |  | Autoinducer Synthesis | 1 | 0,55 |
| PA3476 | *rhlI* |  | Autoinducer Synthesis | -4 | 1,83E-45 |
| PA3477 | *rhlR* |  | Transcriptional Regulation | -3 | 2,39E-29 |

<sup>a</sup>All genes presented are categorized as being QS regulated by Hentzer et al., [22]. The results are based on three independent experiments.

Besides decreasing the expression of genes encoding virulence factors, Z-ethylthio enynone affected an abundant number of adhesion genes. Genes belonging to the *tad* (tight adhesion) loci (**Table S4**) which encode components required for the biosynthesis of the flp pili (type iVb pili) used for adherence to surfaces and twitching motility. The *cup* (chaperone–usher pathway) gene encodes components enabling fimbrial subunit assembly of the type IV pili. In *P. aeruginosa* PA01 four *cup* clusters (*cupA-C,E*) have been identified and characterized. Of the four clusters, only *cupE* is downregulated by Z-ethylthio enynone (**Table S5**). The three *cupA-C* clusters are regulated by c-di-GMP whereas the expression of *cupE* is regulated by the PprAB two-component system which is a part of the *tad* locus (37). The *cupE* gene cluster encodes fimbriae important for early stages of biofilm formation. Two of the three type VI secretion systems were significantly downregulated (H2-T6SS and H3-T6SS) (**Table S6**). Both are positively regulated by the QS system, whereas the H1-T6SS is negatively regulated (38).

The reduction in gene expression is following the same trend when comparing the qRT-PCR results of *lasB*, *rhlA* and *pqsA* with the results of the RNA-seq analysis. The qRT-PCR data showed Z-ethylthio enynone decreased the expression of *rhlA* 18-fold, *lasB* 74-fold and increased *pqsA* 3-fold and the RNA-seq analysis showed that Z-ethylthio enynone decreased the expression of *rhlA* 17-fold, *lasB* 76-fold and increased *pqsA* 3-fold.

### Alkylthio enynone compounds decrease production of QS-regulated virulence factors

Production of two important QS-regulated virulence factors were quantified to investigate if the effect of Z-ethylthio enynone on transcriptional expression of QS genes correspondingly influences protein synthesis. Rhamnolipid encoded by the *rhlA* (PA3479), *rhlB* (PA3478) and *rhlC* (PA1131) genes is controlled by the Rhl and the PQS system. Production is initiated in the early stationary phase of a shaken batch culture. Wild-type (WT) PAO1 cultures were treated with 100 μM of Z/E-ethylthio enynone, Z-ethylthio enynone, and Z/E-propylthio enynone and samples were retrieved at an OD_600_ of 2.0 (stationary growth phase) where rhamnolipid synthesis is fully induced. All three alkylthio enynone compounds significantly decreased the production of rhamnolipid compared to an untreated control **(Fig. 5A)**. Synthesis of the phenazine pyocyanin is firstly triggered by induction of the phenazine operon (*phzA-G*). In addition, seven genes have been reported to be involved in the synthesis of pyocyanin where the *phzM* (PA4209) and *phzS* (PA4217) genes are central. For the measurements of pyocyanin production cultures were grown and samples retrieved as for the experiment assessing the effect of the alkylthio enynone compound on rhamnolipid production. All three alkylthio enynone compounds significantly decreased pyocyanin production **(Fig. 5B)**.

**FIG 5.**
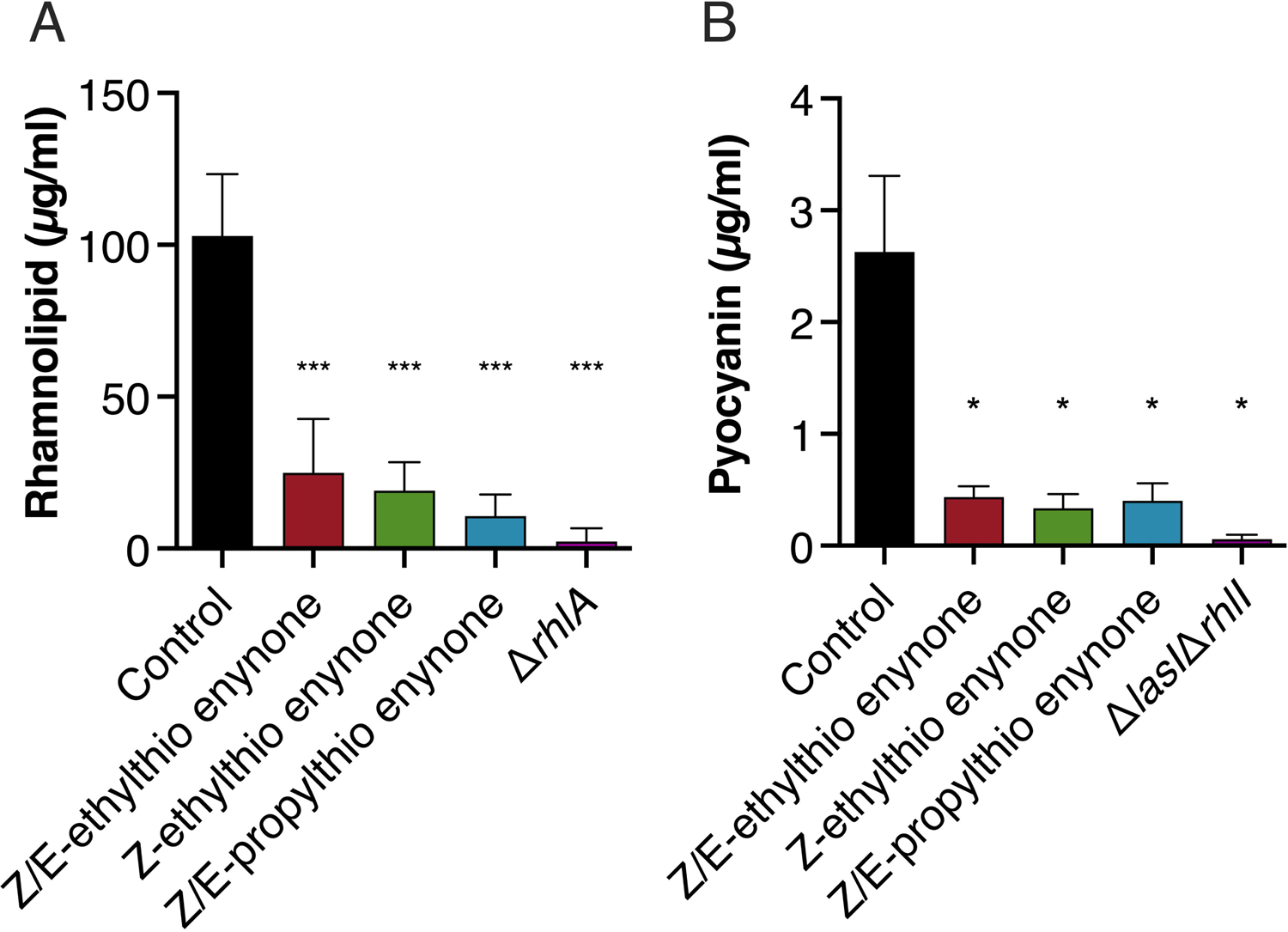
Effect of alkylthio enynone compounds on production of QS controlled virulence factors. A) rhamnolipid and (B) pyocyanin. Concentrations in batch cultures of *P. aeruginosa* grown with 100 μM of Z/E-ethylthio enynone, Z-ethylthio enynone and Z/E-propylthio enynone. Samples were retrieved at OD_600_ of 2.0 and rhamnolipids were quantified using the orcinol assay. Pyocyanin was quantified by extraction into an acidic solution followed by A520 measurements. The results are based on three independent experiments. Error bars represent means ± standard deviation (SD). Significance values between control and treatment are based on unpaired Welch t-test (* p ≤ 0.05, ** p ≤ 0.01 and *** p ≤ 0.001).

### Mutations in *mexEF* and *mexT* genes change the response of *P. aeruginosa* to treatment with Z-ethylthio enynone

To investigate the involvement of potential genes responsible for the QS inhibitory activity of the Z-ethylthio enynone compound, a mutant library was created by transposon mutagenesis in a newly constructed gentamicin sensitive PAO1 CTX-*lasB*-*gfp* reporter strain. The original gentamicin resistant *lasB*-*gfp* monitor strain could not be used as the transposon employed for the library creation required gentamicin for positive selection of the transposon mutants.

The reasoning behind utilizing a fluorescence-based *lasB*-*gfp* reporter strain for the transposon mutagenesis is the enhanced sensitivity which makes it possible to gauge uninterrupted QS activity in real-time. The differences in fluorescence level of colonies are easily detectable as mutants harboring a transposon in a gene that is vital for the activity of Z-ethylthio enynones appear bright, whereas colonies affected by the QS inhibitory activity of the compound appear dim as the transposon is located in a gene unrelated to QS, when monitored with a fluorescence microscope.

The transposon mutagenesis resulted in approximately 60.000 mutants. The mutant colonies exhibiting fluorescence in the presence of 100 μM Z-ethylthio enynone were isolated as these colonies demonstrated unaltered expression of *lasB* despite of the treatment. In total, 56 fluorescent colonies were isolated and all of them were further validated by fluorescence measurements with and without Z-ethylthio enynone treatment in a microplate setup. The region flanking the transposons in ten mutants that demonstrated the highest *lasB* activity in the presence of Z-ethylthio enynone was subsequently sequenced. The identity of the affected genes was mapped by comparing the sequence to the PAO1 genome. Eight mutants all had transposon insertions in the operon encoding the MexEF-OprN efflux-pump, and its transcriptional activator MexT (**Fig. 6A**). Two isolates had mutations that could not be linked to the reference genome of PAO1, and they were therefore discarded for further analysis as the junction was identified to be located in the PlasB-*gfp*(ASV)::Plac-lasR expression cassette. The gene encoding the transcriptional regulator MexT has previously been observed to be a hotspot for mutations affecting the flow generated by the efflux pumps in WT strains. The mutations exist as single-nucleotide polymorphisms (39) and the inclusion or absence of an 8-bp fragment (40, 41) each one of them leading to MexT alteration. To ensure that this mutation is not affecting the results of this study the sequence of the strain was investigated for mutations and no genomic changes of the regulatory parts of the efflux pump were identified.

**FIG 6.**
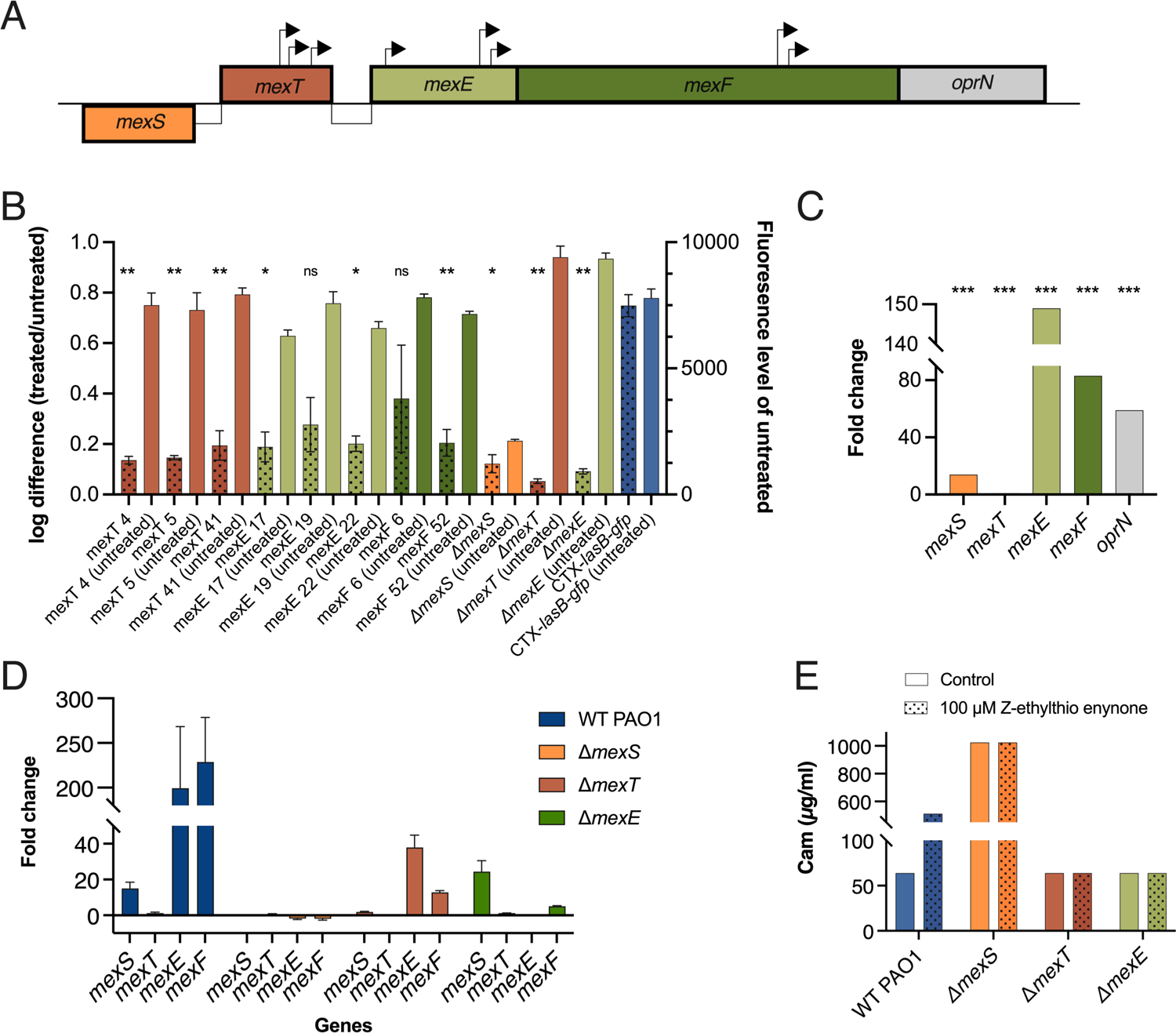
Transposon insertions in *mexT, mexE* and *mexF* genes change the response of *P. aeruginosa* PAO1 to treatment with Z-ethylthio enynone. A) Colonies with transposons in *mexT*, *mexE* and *mexF* isolated from the transposon mutagenesis. Arrows represents the site and the direction of the transposon insertions in the respective gene. B) Effect of 100 μM Z-ethylthio enynone on the fluorescence output of transposons with insertions in *mexT*, *mexE* and *mexF* isolated from the transposon mutagenesis, the constructed *mexS*, *mexT* and *mexE* knockout mutants and the CTX-*lasB-gfp* monitor strain. The bars with pattern show log difference in fluorescence output between treated and untreated. The bars with no pattern show fluorescence level of untreated strains. The results are based on three independent experiments. Error bars represents means ± standard deviation (SD). Significance levels of log difference between treated and untreated control CTX-*lasB-gfp* and the different mutants are based on One-way ANOVA analysis with Dunnett multiple comparisons test, (ns p > 0.05, * p ≤ 0.05, ** p ≤ 0.01 and *** p ≤ 0.001). C) Fold change in gene expression of *mexS*, *mexT* and the *mexEF-oprN* operon measured by RNA-seq analysis of cultures treated with 100 μM Z-ethylthio enynone compared to an untreated control. Statistical methods are based on unpaired t-test, (*** p ≤ 0.001). D) Fold change in gene expression of *mexS*, *mexT, mexE,* and *mexF* in the WT PAO1, Δ*mexS*, Δ*mexT* and Δ*mexE* background strains measured by qRT-PCR analysis of cultures treated with 100 μM Z-ethylthio enynone compared to an untreated control. The results are based on three independent experiments. Error bars represents means ± standard deviation (SD). E) MIC values of chloramphenicol in WT PAO1, *mexS*, *mexT* and *mexE* knockout mutants cultured with and without 100 μM Z-ethylthio enynone. WT PAO1. CAM: chloramphenicol. The results are based on three independent experiments.

Three defined in-frame knockout mutants of *mexS, mexT* and *mexE* were constructed to further investigate the involvement of the MexEF-OprN efflux pump in the observed activity of Z-ethylthio enynone. Mutants with transposons located in the *mexS* gene were not isolated from the library screening, however, this mutant was included as MexS is the main transcriptional repressor of *mexEF-oprN*, and it could provide valuable insight if treatment with the Z-ethylthio enynone compound resulted in a similar phenotype as that of the Δ*mexS* mutant. The fluorescence of the constructed knockout strains was measured when exposed to the Z-ethylthio enynone compound and compared to untreated counterparts to determine if the mode of action of Z-ethylthio enynone is dependent on one or more elements of the *mexEF-oprN* operon (**Fig. 6B+ S5**). Overall, the *mexT* and *mexE* knockout strains undergoing treatment with the Z-ethylthio enynone compound had a slightly increased fluorescence output in contrast to the WT *lasB-gfp* strain. Whereas the untreated *mexS* knockout strain did not show any fluorescence indicating an unfunctional QS system because of a highly expressed MexEF-oprN efflux pump. Except for two strains with transposon insert in *mexE* (19) and in *mexF* (6) all strains with transposons and the *mexT* and *mexE* knockout mutants showed a significant difference when treated as compared to the WT (CTX-lasB-gfp). This indicates very strongly that the MexEF-oprN efflux pump is involved. This data was supported by the transcriptomic analysis (**Fig. 6C**) as genes encoding *mexE* and *mexF* were among the most upregulated in PAO1 WT cultures treated with Z-ethylthio enynone compound. To further explore gene expression of the *mexEF-oprN* efflux pump when treated with Z-ethylthio enynone compound fold change in gene expression of *mexS*, *mexT*, *mexE* and *mexF* between treated and untreated were measured with qRT-PCR in a WT PAO1 and the constructed *mexS*, *mexT* and *mexE* knockout strains. The fold change between treated and untreated WT PAO1 measured by qRT-PCR (**Fig. 6D**) is similar to the results of the transcriptomic analysis showing a significantly increased fold change of *mexS* (14-fold), *mexE* (150-fold) and *mexF* (83-fold) when treated with Z-ethylthio enynone (**Fig. 6C**). For the *mexS* knockout mutant no change in fold change between treated and untreated can be measured. For the *mexT* knockout mutant, there is no change in the gene expression of the *mexS* gene when treated and the fold change for the *mexE* gene and the *mexF* gene is 80% and 95% lower compared to the WT, respectively. For the *mexE* knockout mutant, the fold change of the *mexS* (24-fold) and *mexT* (1.2-fold) genes are very similar to the WT whereas the fold change for the *mexF* gene is 98% lower (**Fig. 6D**).

As one of the primary functions of the efflux pumps is to alleviate the cell from toxic compounds such as the antibiotic chloramphenicol, it was deemed necessary to test the hypothesis that the Z-ethylthio enynone compound stimulates the efflux pumps into generating elevated efflux. As expected, the *mexS* knockout mutant exhibited the highest chloramphenicol MIC value (1024 µg/ml) among all the samples regardless of the presence of Z-ethylthio enynone (**Fig. 6E**). There was no change in MIC values (64 µg/ml) of the *mexT* and *mexE* knockout mutants compared to their untreated counterparts, whereas treatment with Z-ethylthio enynone compound increased the MIC value of the WT PAO1 from 64 µg/ml to 512 µg/ml.

### The QSI compound iberin and Z-ethylthio enynone demonstrates elevated toxicity in the mexEF defective mutants

We have previously documented the QSI capabilities of compounds originating from various food sources (42). Among these compounds were iberin, which is isolated not only from horseradish but also from various other members of the Brassicaceae family. Iberin exhibits distinct QSI-related traits similar to Z-ethylthio enynone, including the ability to reduce QS-controlled virulence factors and possessing nearly identical transcriptomic profiles. To investigate if iberińs QSI ability shares a similar mode of action as Z-ethylthio enynone, specifically regarding the upregulation of the MexEF-oprN efflux pump, we conducted experiments using four CTX-*lasB*-*gfp* reporter strains (WT, Δ*mexS,* Δ*mexT,* Δ*mexE).* These strains were grown in the presence of different concentrations of iberin ranging from 1.5 to 400 µM.

The growth of the *mexT* and *mexE* knockout mutants was significantly inhibited in the presence of iberin in concentrations ranging from 50 to 400 µM and therefore it was not possible to obtain any useable results of *lasB* expression. This observation suggests that the absence of a functional MexEF-oprN efflux pump results in heightened susceptibility to compounds that would otherwise be non-bactericidal in a WT background (**Fig. S6**).

As a continuation to the observations with iberin, the four CTX-*lasB*-*gfp* reporter strains (WT, Δ*mexS,* Δ*mexT,* Δ*mexE)* were grown in the presence of Z-ethylthio enynone in higher concentrations (100 to 800 µM) compared to the rest of the experiments in this study. Due to interference of DMSO with bacterial growth, it was not possible to reach higher concentrations beyond 800 µM. Similar to the findings with iberin, the presence of a non-functional MexEF-oprN efflux pump, as observed in the *mexT* and *mexE* knockout mutants, reduced the tolerance towards the compound resulting in a significantly increased growth inhibition for both strains at 200 µM and higher. The WT PAO1 showed significant growth inhibition at 800 µM Z-ethylthio enynone, whereas the growth of the *mexS* knockout mutant was not affected by any of the tested concentrations. (**Fig. S7**). The growth of the *mexE* and *mexT* knockout mutants was not changed with or without treatment of 100 µM Z-ethylthio enynone (**Fig. S8**).

### Z-ethylthio enynone triggers mexEF facilitated depletion of intracellular QS signal molecules

To investigate if treatment with Z-ethylthio enynone alters the distribution of signal molecules from the cell to the environment, the intra- and extracellular concentrations of AHL signal molecules were measured. Following treatment, the concentration of the signal molecules 3-oxo-C12-HSL and C4-HSL significantly changed compared to an untreated control in a WT strain. The extracellular concentration of 3-oxo-C12-HSL in the WT strain, increased as a response to the Z-ethylthio enynone treatment (*** p≤ 0.001), and the corresponding intracellular concentration was decreased (*** p≤ 0.001) (**Fig. 7A**). Both the intra- and extracellular level of C4-HSL were significantly reduced, * P ≤ 0.05 and ** p≤ 0.01, respectively (**Fig. 7B**). For the three knockout mutants (Δ*mexS*, Δ*mexT*, and Δ*mexE*) no difference could be measured between control and treated samples regarding the extracellular and intracellular concentration of 3-oxo-C12-HSL (**Fig. 7A**) and C4-HSL (**Fig. 7B**). For the untreated MexS knockout mutant were the intracellular concentration of 3-oxo-C12-HSL significantly lower (** p≤ 0.01) than for the WT PAO1, whereas there was no significant difference between the two strains when treated and they showed a very similar concentration of signal molecule. For the WT PAO1 treated and untreated there was a significant difference (* P ≤ 0.05) to both the MexE and MexT knockout mutants with a 2-fold decrease in concentration for the untreated and a 12-fold increase for the treated Mex mutants (**Fig. 7C**).

**FIG 7.**
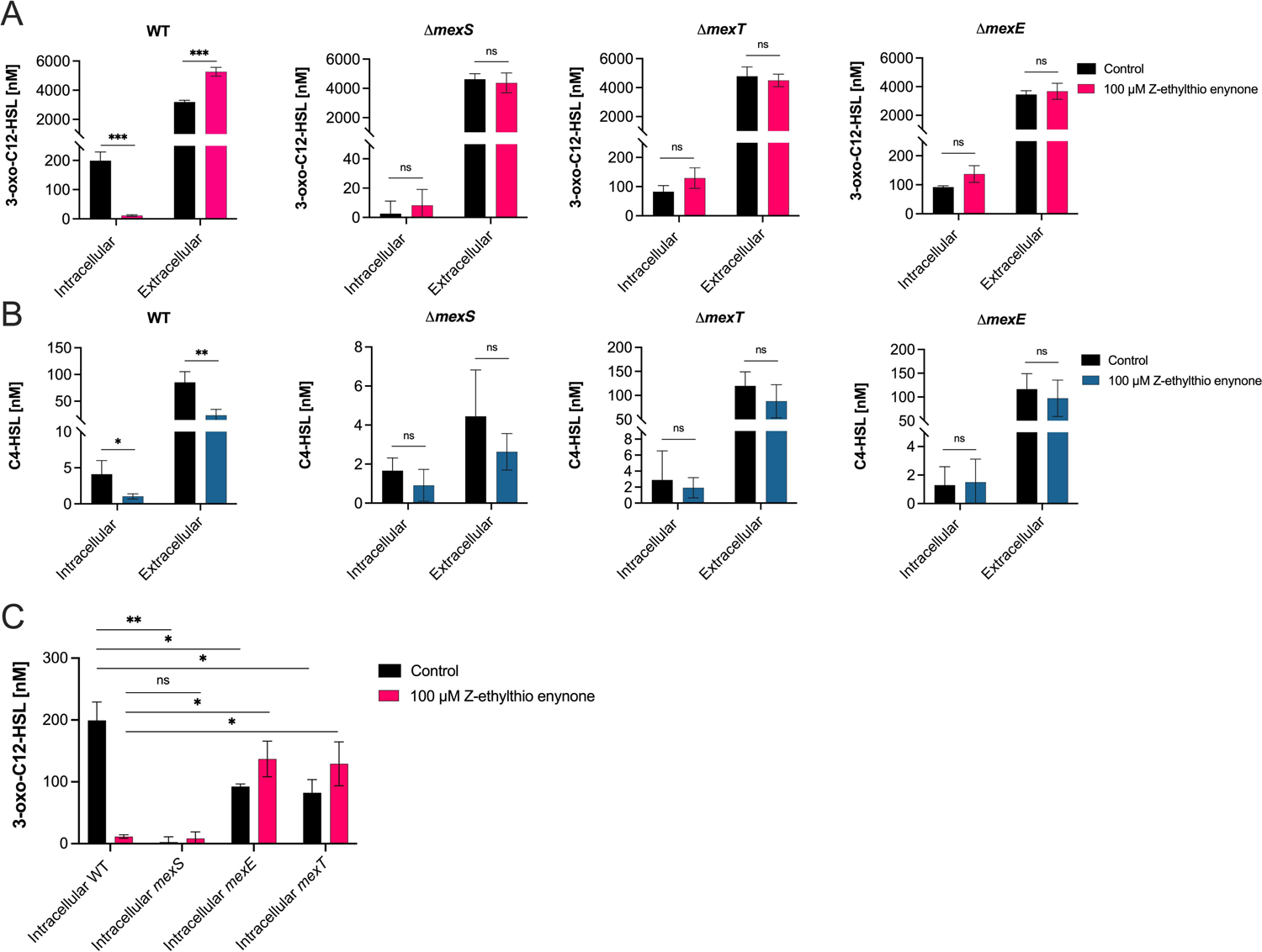
Quantification of 3-oxo-C12-HSL (A) and C4-HSL (B) QS signal molecules in WT PAO1 and the *mexS*, *mexT* and *mexE* knockout mutants treated without and with 100 μM Z-ethylthio enynone. The concentration is measured both intra- and extracellular. Intracellular levels of 3-oxo-C12-HSL (C) compared between the *mexS*, *mexT* and *mexE* knockout mutants and the WT PAO1. The results are based on three independent experiments normalized to the total concentration of signal molecules for each of the two signal molecules. Error bars represent means ± standard deviation (SD). Significance values between control and treatment are based on unpaired Welch t-test (ns p > 0.05, * p ≤ 0.05, ** p ≤ 0.01 and *** p ≤ 0.001).

## DISCUSSION

A large proportion of hospital-acquired infections are caused by bacterial biofilms (43, 44). Due to the ability of biofilms to tolerate high doses of antibiotics, they are a common cause of persistent infections, hospitalizations, patient suffering, and reduced quality of life. Since our current treatment regimens in many cases fail to cure biofilm-based infections, new anti-biofilm drugs and novel treatment strategies are urgently needed. The critical need for the development of new antimicrobial products against *P. aeruginosa* infection is emphasized by the high frequency of patients with reoccurring infections that often respond poorly to antibiotic treatments (45).

In this study, we present three chemical compounds discovered to be potent QSIs using high-throughput screening. The mechanism of action was identified to be dependent on the MexEF-oprN efflux pump using transposon mutagenesis on a newly constructed PAO1 CTX-*lasB-gfp* reporter strain, and the use of different Mex knockout mutants.

The identified compounds in this study are strong QSIs decreasing the expression of a high number of QS genes as well as decreasing the production of the QS-regulated virulence factors, rhamnolipid and pyocyanin. The phenotypes of the selected transposon mutants together with the constructed *mexE* and *mexT* knockout mutants and the highly increased expression of *mexE* (150-fold) *mexF* (83-fold) and *oprN* (59-fold) indicate that the MexEF-oprN efflux pump plays an important role in alkylthio enynones mechanism of action. The increased expression of *mexS* (14-fold) together with the status quo for *mexT* expression does not correspond with studies showing that *mexS* has a negative effect on *mexT* expression (27). In the contrary, studies have shown by gene expression analysis that a functional *mexS* and *mexT* are required for activation of *mexEF-oprN* and increased expression of *mexS,* and an unchanged expression of *mexT* can increase the expression of *mexE* (23, 46, 47) which is similar to the results of the transcriptome of alkylthio enynone treated PAO1 presented in this study. In addition, an increase in expression of *mexEF-oprN* (12.9-fold) and *mexS* (13.3-fold) has been shown to occur when *P. aeruginosa* PAO1 interact with airway epithelial cells (48) further indicating that an increase in *mexS* expression can lead to an increase in *mexEF-oprN* expression and not necessarily results in a decrease in expression of the efflux pump. This suggests that there are alternative regulatory mechanisms of MexS and MexT regulation of the MexEF-oprN efflux pump, which are not fully elucidated.

The role of the MexEF-oprN efflux pump has primarily been linked to aiding the antibiotic tolerance of *P. aeruginosa*, yet growing evidence points towards that this function is not solely limited to antibiotics but rather as a mechanism to relieve the cell from various forms of stress (49, 50). While we can only speculate on the exact chemical properties of the alkylthio enynone compound, we hypothesize that the stimulation of the efflux pumps stems from the compound or degradation parts generating a stress response, as it was observed that treated strains with a defective MexEF-oprN efflux pump had a decrease in growth compared to a WT PAO1. A similar trend was observed with QS inhibitor iberin. Additionally, there was a very similar pattern of downregulated QS-regulated genes between Z-ethylthio enynone and iberin-treated *P. aeruginosa* cultures and more importantly, the expression of genes associated with the MexEF-oprN efflux pump (*mexS, mexT,* and *mexEF-oprN*) were up or downregulated similarly when exposed to iberin or alkylthio enynone (**Table S8**). Together with the increase in growth inhibitory activity of MexEF-oprN defective strains by these two compounds indicate that they share a similar mechanism of action. Additional reasoning behind this comparison of mechanisms of action between these two QSI compounds originates from research on how *P. aeruginosa* utilizes the MexEF-oprN efflux pump to combat oxidative stress from various chemical sources such as nitric- and superoxide (51), disulfide (52), and reactive oxygen species(53). When comparing these data to our transcriptomic analysis of PAO1 treated with the Z-ethylthio enynone compound, a similar pattern in the stress-induced activation of the genes associated with oxidative stress and the treatment with Z-ethylthio enynone is observed. Among these genes was *xenB* that encodes an enzyme responsible for nitration and thereby aiding the bacteria in reducing nitrosative stress that occurs during treatment with chloramphenicol. Not only is *xenB* upregulated 10-fold in our case, but several genes highlighted in these studies are also upregulated as a response to Z-ethylthio enynone treatment (PA1744, PA1970, PA2486, PA2759, PA2811, PA2812, PA2813, PA3229, *xenB*, PA4623, PA4881). This includes having 3 candidates in the top 5 of the most upregulated genes in response to Z-ethylthio enynone treatment (**Table S7**) which are all connected with the MexEF-oprN efflux pump as previously reported by (47). However, the MexEF-oprN efflux pump is not the sole defense against oxidative stress, PAO1, and other bacterial species also employ antioxidative enzymes synthesized by the core regulator *oxyR* (54). Neither *OxyR* nor regulatory-connected genes were affected by the addition of Z-ethylthio enynone. No mutants with genes not related to the MexEF-oprN efflux pump were identified with the transposon mutagenesis, which further confirms that Z-ethylthio enynone exclusively depends on the MexEF-oprN efflux pump for the observed QSI effect.

There is no indication that the observed QS inhibitory activity of the Z-ethylthio enynone compound is caused by directly interacting with the core regulatory elements of the QS systems. Our data suggests that the cause of this phenotypic change stems from the QS system being prevented from reaching the required concentration threshold by active drainage of intracellular QS signal molecules through activation of the MexEF-oprN efflux pump. The intracellular concentration of 3-oxo-C12-HSL was significantly reduced (*** p≤ 0.001) in the WT, while the extracellular concentration was likewise significantly increased (*** p≤ 0.001) upon treatment with Z-ethylthio enynone (**Fig. 7A**). The Rhl system of the WT PAO1 was correspondingly affected by the treatment with both the intracellular and extracellular concentration of C4-HSL (**Fig. 7B**) being significantly reduced, (* p≤ 0.05) and (** p≤ 0.01), respectively. This is as expected as the QS system in *P. aeruginosa* is organized in a hierarchy-like structure where a negatively affected Las-system will delay the activation of the Rhl system. We, therefore, believe that the observed inhibition of the QS-system originates from the forceful relocation of intracellular signal molecules belonging to the Las-system, as it is being actively pumped out of the cell by the MexEF-oprN efflux pump, thereby leading to the elevated concentration of signal molecules located in the surrounding environment. The efflux thereby prevents 3-oxo-C12-HSL from accumulating in the cell as the passive diffusion fails to counteract this process, thus blocking the potential of reaching the required concentration threshold and thereby delaying the synthesis of signal molecules related to the RHL system.

Additionally, this correlates with the observed inhibition of our QS-reporter strains, the reduction in synthesis of the QS-controlled virulence factors, pyocyanin, and rhamnolipid (**Fig. 5A+B**) as well as the greatly decreased gene expression of the corresponding QS-related genes in response to the treatment (**Fig. 3E**). The effect of signal molecule efflux on QS is even more evident in the *mexS* knockout mutant with its persistent efflux regardless of treatment. This had a comparable effect on the distribution of the signal molecules to that of the treated WT strain albeit with much lower concentrations of both signal molecules. Similar has been documented previously by (55).

It is interesting however that the *mexS* knockout mutant can disrupt the activation of the Rhl system to such an extreme degree compared to that of the treated WT PAO1. A possible explanation for this could lie in the difference in velocity of efflux caused by the MexEF-oprN efflux pump. While *mexE* and *mexF* were among the most upregulated genes in the transcriptomic analysis of the WT strain treated with Z-ethylthio enynone, 149 and 83-fold, respectively (**Fig. 6D**), it has previously been shown that the expression of these two genes in an nfxC background was upregulated to a staggering 2033 and 1382-fold, respectively (27). Indicating there could be a correlation between gene expression and the velocity of efflux. This trend corresponds to our other experiments, i) Δ*mexS* was the only strain to be largely unaffected by the elevated concentrations of Z-ethylthio enynone and iberin (**Fig. S6+S7**), ii) Δ*mexS* had a MIC value of CAM at 1024 µg/mL, which is two-fold higher than the treated WT PAO1 (**Fig. 6D**), and iii) the previously discussed difference in AHL content (**Fig. 7A+B**).

It could be theorized that *mexT* and *mexE* knockout mutants with deficient MexEF-oprN efflux pumps result in an accumulation of intracellular signal molecules. While we have shown that the MexEF-oprN efflux pump can transport signal molecules, it is also well documented that AHL molecules are not the main substrates of these pumps as well as gram-negative bacteria being known to depend on passive diffusion to transport the signal molecules. It is seen that *mexT* and *mexE* knockout mutants have a functional QS system, as there is no significant difference between the control and treatment in regards to the concentration or location of the AHL (**Fig. 7A+B**).

We conclude that the expulsion of QS-related signal molecules by the MexEF-oprN efflux pumps is thus a necessary sacrifice for the cell to avoid the cellular stress of Z-ethylthio enynone at the cost of losing control of the QS system.

One could argue that the use of the previous version of the PAO1-*lasB* monitor strain in the initial chemical screening had an unwanted bias towards compounds utilizing the MexEF-oprN pump for its QSI capabilities, however, we have previously discovered the potent QSI ajoene using similar quantification methods, which only affect a very limited set of genes by relying on interaction with small regulatory RNA rather than generating an efflux response (34, 56). Notably, we were unable to isolate any mutants harboring a mutation in the *mexS* gene, most likely because a non-functional MexS is identical to the *nfxC* phenotype. The phenotypic characterization of the nfxC phenotype was first described by (57), after PAO1 was exposed to fluoroquinolones it developed cross-resistance tolerance towards multiple classes of antibiotics because of inactivating the MexEF repressor MexS by mutation and thereby creating enhanced efflux (25). This phenotypic change most likely explains why we could not isolate any *mexS* mutants, as a transposon insertion that interrupts the function of *mexS* would upregulate the transcription factor MexT thereby generating efflux and the expulsion of QS signal molecules, which results in a missing GFP signal from the CTX-*lasB-gfp* monitor strain. Besides that, we were able to isolate mutants that exclusively had a mutation in one of the three genes responsible for the regulation and assembly of the MexEF-oprN efflux pump, with *mexT* and *mexE* both having 3 unique transposon inserts each, and *mexF* had two transposon inserts in the same gene emphasizing that the new CTX-*lasB-gfp* monitor strain is sensitive enough to gauge QSI activity *in vitro* and being applied to identify possible QS related drug targets by the use of the novel transposon mutagenesis setup presented in this study. Possible mechanisms of action are presented in **Fig. 8**.

**FIG 8.**
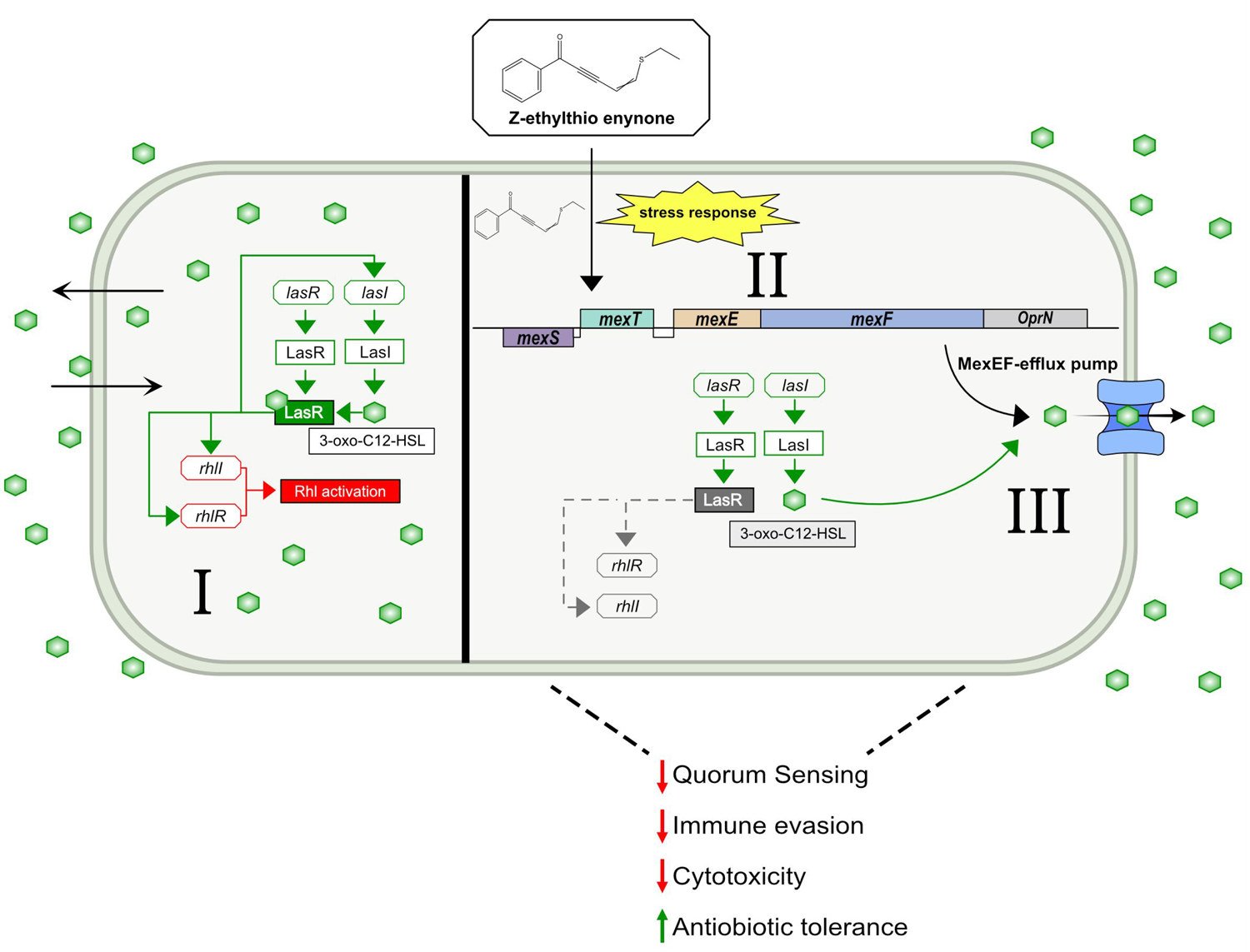
The proposed mechanism of action for alkylthio enynone. **I**) The uninterrupted QS system of *P. aeruginosa* with freely diffusible signal molecule 3-oxo-C12-HSL which is synthesized by LasI. When the concentration threshold has been reached, the signal molecule binds to its receptor LasR, which leads to a positive feedback loop increasing the expression of *lasI* and thereby enhancing the synthesis of additional 3-oxo-C12-HSL, and initiation of synthesis of the Rhl system. **II**) Both the alkyne and the alkene are reactive towards alkylthiols. **III**) Alkylthio enynone or a chemical byproduct prompts a stress response thus promoting the expression of genes related to the MexEF-oprN efflux pump to alleviate stress from the cell. **IV**) 3-oxo-C12-HSL is synthesized as in **I**) but as the consequence of the elevated efflux motivated by the stress response, the cell is depleted of the intracellular signal molecules and in this manner, the cell is unable to reach the concentration threshold of the signal molecule and thus reduced synthesis of QS controlled virulence factors but also elevated tolerance towards antibiotics.

Further work must be done to confirm if other QSI compounds share similar action by implementing the newly constructed CTX-*lasB-gfp* reporter strain in conjunction with transposon mutagenesis for localizing QSI target as described in this paper. It should be possible to revalidate the former generations of QSIs as well as future compounds thereby getting deeper insight into the QS system of *P. aeruginosa* and possibly future treatment options to combat chronic infections.

## MATERIALS AND METHODS

### Bacterial strains and cultivation

Overnight (ON) cultures were grown in LB media at 37°C with shaking (200 rpm) and supplemented with antibiotics where appropriate (all strains used are described in **Table S1**). B minimal media [48] with 10% A-10 (58) supplemented with 0.5% (wt/vol) casamino acid and 0.5% (wt/vol) glucose (ABGC media) was used in the experiments described in the following sections if not stated otherwise.

### Initial library screens for quorum sensing inhibition

The chemical library consists of 3280 compounds divided into two collections: Analyticon (2080 compounds) and Phytoquest (1200 compounds). The 3280 compounds were tested in a final concentration of 10 µM with the *rhlA-gfp* [30] reporter strain in 96 wells microtiter dish (Black Isoplate^®^, Perkin Elmer). The cells were added in a final concentration of OD_600_=0.1 and final volume of 200 µl in ABGC medium and incubated at 37°C with shaken (200 rpm). Growth (OD_450_) and fluorescence (GFP expression (excitation wavelength, 485 nm and emission wavelength, 535 nm)) was measured after 12 hours using a Victor plate reader (Perkin Elmer). Eighty-nine compounds showing activity were re-tested with the *lasB-gfp* (31) and *rhlA-gfp* (30) monitor strains. The 10 most active compounds were selected, and new batches of the compounds were tested in two-fold serial dilutions starting from a concentration of 100 µM. The 10 compounds were tested with the *lasB-gfp*, *rhlA-gfp*, *rsmY-gfp* (34) and *rsmZ-gfp* (34) monitor strains (see determination of IC50 values using QSI screen).

### Chemically synthesized alkylthio enynone compounds

The resynthesized alkylthio enynone compounds were prepared as 100 mM stock dissolved in dimethyl sulfoxide (DMSO) and stored at −20°C (see Supplemental Material for a specified description of synthetic procedures).

### Growth curve

A PAO1 wildtype (WT) ON culture was diluted to OD_600_ of 0.01 in conical flasks containing ABGC media, each of the flasks were then supplemented with one of the alkylthio enynone analogs, resulting in a final concentration of 100 µM. Equal volumes of DMSO were added to the control culture. The conical flasks were incubated at 37°C with shaking (200 rpm). Growth was measured as OD_600_ every hour for the first two hours and every 30 min. thereafter, for a total run time of 8 hours.

### Determination of IC50 values using quorum sensing inhibitory screening

The reporter systems *lasB*-*gfp* (31), *rhlA*-*gfp* (30), *pqsA*-*gfp* (59), *rsmY*-*gfp* (34) and *rsm*Z-*gfp* (34)) were used to determine an IC_50_ (half maximum inhibitory concentration) value in relation to the QSI capabilities of the alkylthio enynone compounds. A concentration gradient of the compounds was made by two-fold dilution using fresh ABGC media supplemented with 0.4% DMSO in 96 wells microtiter dish (Black Isoplate^®^, Perkin Elmer). The monitor strains were diluted to OD_450_ of 0.1 and transferred to the wells, resulting in a final test volume of 200 µL. An Infinite F200 pro (TECAN^TM^) plate reader was used to monitor growth at OD_450_, and GFP activity was measured as fluorescence at an excitation wavelength at 485 nm and emission wavelength at 535 nm every 15 min. for 18 hours. The temperature was held constant at 34°C.

To Calculate the IC50 value of the alkylthio enynone compounds the slopes of the relative fluorescent unit/OD_450_ ratio measurements between two time points (*lasB*: 220-360 min, *rhlA*: 200-320 min) were used to generate dose-response curves in Graphpad Prism 8.

### Cell harvest for RNA purification

A PAO1 WT ON culture was diluted to OD_600_ of 0.01 in a conical flask containing 100 ml of AB media supplemented with 0.5% casamino acid and incubated at 37°C with shaken (200 rpm). At OD_600_ of 0.5 the culture was diluted to an OD_600_ of 0.1 and incubated again under the same condition. At an OD_600_ of 0.5 (exponential growth), the culture was split into separated conical flasks containing 20 ml of the culture and supplemented with one of the three alkylthio enynone compounds to a final concentration of 100 μM. The untreated cultures were supplied with the same volume of DMSO. At OD_600_ of 2.0, the bacterial cultures were harvested and RNAprotect Bacteria Reagent (QIAGEN) was added in a ratio of 2:1, and the accompanying quick start protocol was followed. The pellet was stored at −80°C until RNA extraction.

### RNA extraction

RNA extraction was performed using RNeasy Mini Kit (Qiagen) according to the manufacturers’ instructions with DNase 1 (Biolabs) treatment. Total RNA yield and possible DNA contamination were evaluated using a Nanodrop spectrophotometer (Thermo Scientific) and possible fragmentation of the RNA was spotted on a 1% agarose gel. The integrity of the RNA was tested on a Bioanalyzer 2100 using the Prokaryote Total RNA Nano kit (Agilent Technologies) and RNA Integrity Numbers (RIN) over 7 was accepted for RNA sequencing (RNA-seq) and quantitative Real Time-PCR (qRT-PCR).

### cDNA synthesis

cDNA was synthesized from 1 μg of RNA by using a High-capacity RNA-to-cDNA master mix (Applied Biosystems™) in a total reaction volume of 20 µl. A negative control (NoRT) without reverse transcriptase added was included in every run. The reaction was done using a preset program, running at 37°C for 60 min. and 95°C for 5 min, and the synthesized cDNA was then stored at −20°C if not used immediately for qRT-PCR.

### Quantitative Real Time-PCR

Primers were ordered from TAG Copenhagen with a synthesis scale of 0.04 μmol and purified with RP-column (sequence of all primers are presented in **Table S2**). The metabolic gene *rpoD* was used as a reference gene. cDNA was diluted 1:100 and primers were used in a final concentration of 200 nM. qRT-PCR amplification was performed with Power Sybr green master mix (Applied Biosystems™) in a Step One Plus Thermal Cycler (Applied Biosystems). The PCR starts with 95°C for 10 min. followed by forty cycles of denaturation at 95°C for 15s, annealing/extension at 60°C for 60s.

### RNA-sequencing and data analysis

The isolated RNA was shipped to Beijing Genomic Institute (BGI, Hong Kong) on dry ice for rRNA depletion, library construction, and sequencing. Reads were then aligned to the *P. aeruginosa* reference genome (www.pseudomonas.com, assembly: GCF_000006765.1, RefSeq) using STAR v2.6.1a (60). All reads mapping to protein-coding gene features (CDS) were counted using the ‘quantMode GeneCounts’ option in STAR. Differential gene expression analysis was then performed with DESeq2 v1.16 with default settings (61). The log2 fold-change in normalized gene expression is represented as log2(Normalized Expression_Treated_/Normalized Expression_Control_).

### Measurements of rhamnolipid production

A PAO1 WT and a Δ*rhlA* mutant (62) ON culture were diluted to an OD_600_ of 0.01 in four separate conical flasks containing 20 ml ABGC media. Cultures were treated with 100 μM of the three test compounds, while an equal amount of DMSO was added to the control samples. Cultures were grown for 24 hours before harvest. 1.8 ml cell cultures were harvested by centrifugation at 15.000g for 3 min. The supernatants were collected and filtered through a 0.22 μM syringe filter. 300 μL of filtered cell culture was mixed with 1 mL of ethyl acetate and immediately mixed for 30 seconds followed by centrifugation for 60s at 10000g. 800 μL of the organic phase was transferred to a collecting tube, while the water phase was mixed with 1 ml of ethyl acetate. The ethyl acetate extract was evaporated, and dried samples were dissolved in 300 μL Milli Q water and stored at −20°C until quantification. 100 μL of dissolved extract was mixed with 100 μL 1.6% orcinol and 800 μL 60% H_2_SO_4_ and heated to 80°C for 30 minutes. After cooling the samples, the rhamnolipid content was quantified at OD_420_ in a Synergy 4 Multi-Mode Microplate Reader (BioTek). To calculate the concentration of rhamnolipid a standard curve of rhamnose was made from the following concentrations (0, 0.1, 1, 10, 100 μg/ml). Conversion from rhamnose to rhamnolipid was done by multiplying the values with a factor of 2.5, as it assumed that 1 mg of rhamnose corresponds to 2.5 mg of rhamnolipid (63).

### Measurements of pyocyanin production

ON cultures of a WT PAO1 and a Δ*lasI*Δ*rhlI* double mutant (64) were diluted to an OD_600_ of 0.01 in four separate conical flasks containing 20 ml ABGC media. Treatment, growth, and filtration of the supernatant were performed as described in the section ‘*Measurements of rhamnolipid production’*. Chloroform was added to the supernatant in a 1:2 ratio and vortexed 10 times for 2s. Samples were centrifuged for 10 minutes at 10000g, and the bottom chloroform layer was transferred to a new tube. 0.2M HCL was mixed with the chloroform layer in a 1:2 ratio, and the tube was vortexed 10 times for 2s. The samples were centrifuged at 10000g for 2 min and the pink top layer was transferred to a cuvette for content quantification. The spectrophotometric measurements were done at OD_520_, with 0.2M HCL used as blank. The value was multiplied by 17.072 to get the pyocyanin concentration in µg/ml (65).

### Construction of a gentamicin sensitive PAO1 GTX-*lasB-gfp* reporter

To enable the identification of the gene product(s) targeted by our lead compound Z-alkylthio enynone, we created a gentamicin sensitive PAO1 *lasB-gfp* reporter strain that could serve as the host strain of a pBT20 (66) based transposon library. Initially, the P*_lasB_*-*gfp*(ASV)::P*_lac_*-*lasR* expression cassette of pMHLAS (31) was isolated on a 3,1 kb *Not*I-fragment and inserted into the unique *NotI* site of the chromosomal integration vector mini-CTX2 (67) to give either plasmid pRK1 or pRK2. In pRK1 the *gfp*(ASV) gene and the mini-CTX2 encoded integrase gene is transcribed in opposite directions, whereas the *gfp*(ASV) gene and the integrase gene are transcribed in the same direction in pRK2. Next the P*_lasB_*-*gfp*(ASV)::P*_lac_*-*lasR* expression cassette of pRK2 was inserted into the chromosome of *P. aeruginosa* PAO1 using the 3-step protocol described by Andersen and co-workers (68). Briefly, in step one, a PAO1 transconjugant with plasmid pRK2 inserted into the chromosomal ϕCTX *attB* site was constructed by three-parental mating using *E. coli* DH5-α/pRK2 as donor, *E. coli* HB101/pRK600 as helper, and *P. aeruginosa* PAO1 as recipient. In step two, the tetracycline resistant PAO1 transconjugant was transformed with plasmid pFLP2 encoding Flp recombinase (67) to obtain a pFLP2 transformant where the Fret flanked plasmid backbone of pRK2 had been excised. In step three, the PAO1 transformant containing one chromosomal copy of the P*_lasB_*-*gfp*(ASV)::P*_lac_*-*lasR* expression cassette (from pRK2) was cured for plasmid pFLP2 using sucrose selection, and the resulting gentamicin sensitive PAO1 based *lasB-gfp* reporter strain was designated *CTX-lasB-gfp*. Finally, the chromosomal location of the P*_lasB_*-*gfp*(ASV)::P*_lac_*-*lasR* expression cassette in strain RIKIL1 was verified by PCR using the primer pair Pser-up/Pser-down (67).

### Transposon mutagenesis

The donor strain *E. coli* S17pir/pBT20 was grown on an LB plate supplemented with carbenicillin (100 µg/ml), while the recipient strain PAO1 chromosomal *lasB-gfp* (ASV) was grown on plain LB plates. Both plates were incubated at 37°C ON. The following day both strains were re-streaked on a new set of similar plates. The donor strain was incubated at 37°C ON, while the recipient strain was incubated at 42°C for 16 hours. The bacteria were then scraped off the plate and the OD was adjusted to OD_600_ of 40 for donor and OD_600_ of 20 for recipient. The two strains were then mixed in a 1:1 ratio, spotted on dry LB plates, and incubated for 2 hours at 37°C. The spots were then resuspended in 0.9% saline and a dilution series was plated out on large selective medium plates without alkylthio enynone compound (1,5 % noble agar, B 100, 10% A-10, 100 µM IPTG, 10 mM Citrate and 60 µg/ml GM) to calculate the conjugation efficiency. The remaining cell suspension was stored at 4°C until the following day. The target CFU should be 1500-2000 CFU per plate on large selective media plates containing 100 µM Z-ethylthio enynone. The selective plates were incubated at 37°C for a maximum 18 hours, and colonies still emitting fluorescence were isolated and retested to verify fluorescence, before being stored at −80°C. A random isolated colony that did not demonstrate fluorescence, were likewise isolated and stored at −80°C. The isolated and stored colonies were tested for fluorescence level when treated with 100 µM Z-ethylthio enynone by using the method described in “*Determination of IC50 values using QSI screen*”.

### Arbitrary PCR and sequencing

The template chromosomal DNA preparation and the following PCR reactions were performed as described in (66) with the only deviation from the protocol being the use of premade PCR mastermix. Promega Wizard kit was used to clean the second-round PCR products before being sent for sequencing at Eurofins Genomics. Gene identification of the sequences was done using Blastn against *P. aeruginosa* (taxid:287). The two isolates with a transposon insert junction that could not be mapped to the *P. aeruginosa* refence genome, were discarded for further analysis as the junction was identified to be located in the PlasB-*gfp*(ASV)::Plac-lasR expression cassette by using its plasmid sequence as the reference genome.

### Construction of *mex* deletion mutants

Allelic exchange vectors for introduction of defined in-frame deletions in the mex genes were constructed using pDONRPEX18Gm essentially as previously described (69). The construction of the *mex* deletion PAO1 mutants was performed as described in *“Construction of a gentamicin sensitive PAO1 GTX-lasB-gfp reporter”*. Except for the donor strain being the MS690 cells harboring three different mex allelic exchange vectors, and the antibiotic marker being GM instead of Tet. Each of the three knockout strains was negatively selected on 10% sucrose plates, cured of their antibiotic marker, and validated by running PCR productions on a 0.8% agarose gel, any isolates with a band size above 1kb were discarded with the rest were put into storage.

### Construction of Δ*mexS*/Δ*mexT*/Δ*mexE* GTX-*lasB-gfp* monitors

The sucrose^R^, Gm^S^ knockout mutants were then subjected to the same procedure as described in “*Construction of a gentamicin sensitive PAO1 GTX-lasB-gfp reporter*” to construct knockout mutants encoding the *lasB-gfp* reporter. Flp-mediated excision of the plasmid backbone which includes the Tet^r^ marker was achieved by electroporation of plasmid pFLP2 into each of the knockout mutants and selecting firstly for colonies demonstrating Tet^s^ Cb^r^, and secondly sucrose resistance on LB plates containing no salt and 10% sucrose. The Tet cured recombinants were then frozen. To ensure that the knockout mutants processed intact *mex* genes besides the gene knockout, an additional PCR reaction with the primers covering the genes *mexS* to *mexT*, and *mexT* to *mexE* was conducted. The genes were deemed intact as a fragment size of 2kb appeared for all three knockouts mutants.

### Testing for alkylthio enynone inhibition in the mex-knockouts

The experiment was performed as described in the section “*Determination of IC50 values using quorum sensing inhibitor screen*”.

### Susceptibility to chloramphenicol

To ensure that the alkylthio enynone compound would be able to affect the cells before encountering the antibiotic, cultures were treated with alkylthio enynone or equal concentration of DMSO prior to antibiotic treatment. The alkylthio enynone pretreatment was performed by diluting an ON culture of WT and the three mex knockout mutants to OD_600_ of 0.1 in ABCG media containing 100 µM Z-ethylthio enynone grown at 37°C with shaking (200 rpm) until an OD_600_ of 0.5 was achieved. The pretreated cultures were then diluted to a final OD of 0.1 in fresh media containing alkylthio enynone. A chloramphenicol concentration gradient spanning from 1024 µg/ml to 16 µg/ml was made in a 96-well F-bottom plate (TPP), in which the diluted cultures were added. The cultures were grown at 37°C with shaking (50 rpm) ON, and OD values was recorded the following day on a Victor^TM^ X4 multilabel plate reader (Perkin Elmer).

### Assessment of iberins QSI capability in the MexEF knockout mutants

The experiment was performed as described in the section “*Determination of IC50 values using quorum sensing inhibitor screen”*, with the maximum concentration of iberin being 64 µg/ml (400 µM).

### Quantification of intra- and extracellular signal molecules

Extraction of 3-oxo-C12-HSL and C4-HSL signal molecules was performed as described by (70). An ON culture of WT and each of the mex knockout mutants were diluted to OD_600_ of 0,01 and grown for 20 hours in the presence of 100 µM Z-ethylthio enynone or an equal amount of DMSO acting as the experiment control. Cells were centrifuged and the resulting supernatant was sterile filtered, while the cell pellet was saved for intracellular content analysis. The cell pellet was washed twice in 0,9% saline solution and lysed by the use of methanol, followed by sterile filtration, and left to evaporate. For the extracellular signal molecules, liquid-to-liquid extraction was done using acidified ethyl acetate, and the extract was left to evaporate. Both types of extracts were reconstituted in equal amounts of methanol and stored at −20°C until analysis.

The extracts were transferred to a 96-well microtiter dish (Black Isoplate^®^, Perkin Elmer) and left to evaporate. However, it should be noted that the transferred volume of the extract depends on the QS system and between the intra- and extracellular location of the signal molecule. Additionally, a concentration standard of each of the two-signal molecule was made with recently purchased chemical stocks of 3-oxo-C12-HSL and C4-HSL and fresh ABGC media, the content was then transferred to the microtiter plate. ON culture of one of the two QS-molecule bioreporters (MH 205 and MH155) was diluted to OD_600_ of 0.1 and transferred to the dry wells as well as the standard concentration gradient. The measurement of the GFP signal was performed as described in the section “Determination of IC50 values using quorum sensing inhibitor screen. The signal molecule concentration of the extracts was estimated by comparing its maximum RFU peak to those of the concentration standard, and values were normalized to the total concentration of signal molecules in each sample to take account of the difference in volumes of extract used.

### Statistical Analysis

Statistical significance was based on One-way ANOVA analysis with Dunnett multiple comparisons test for evaluating log difference between treated/untreated mutant strains and WT PAO1, unpaired Welch t-test for evaluating treated versus untreated cultures, and one sample t-test for evaluating significance levels from zero. For calculating p-values the statistical program GraphPad Prism (GraphPad software, Inc., San Diego, USA, p-values ≤ 0.05 were considered significant) was used.

## ACKNOWLEDGEMENTS

This work was supported by grants to MG and TTN from the Independent Research Fund Denmark, the Lundbeck Foundation, and the Danish Ministry of Higher Education and Science (the DK-Openscreen program) as well as a grant to KQ from the Carlsberg Foundation Young Researcher Fellowship (grant No. CF18-0631). Thanks to LifeArc (London, UK) for delivery of compound libraries used in this study.

